# Rice bacterial blight resistance in Burkina Faso through genome editing: Evaluating pathogen and agro-morphological compatibility of genome-edited elite rice varieties

**DOI:** 10.64898/2026.06.24.734420

**Authors:** Soumana Kone, Abdourasmane Kadougoudiou Konate, Antoine Barro, Wolf B. Frommer, Boris Szurek, Eliza P.I. Loo, Issa Wonni

## Abstract

Bacterial leaf blight (BB), caused by *Xanthomonas oryzae* pv. *oryzae* (Xoo), causes yield losses exceeding 50% in affected areas, including the Bagré rice plain in Burkina Faso. Genome-edited (GE’d) rice lines have been successful in tackling BB. Modifications in the Xoo virulence protein target site upstream of three *SWEET* susceptibility genes in two elite rice varieties, IR64 and Ciherang-Sub1, have been demonstrated to confer broad-spectrum resistance to Asian and East African Xoo strains. Here, we evaluate the potential of the GE’d lines as a solution for BB management in Burkina Faso. We challenged the GE’d lines against five locally collected Burkinabè Xoo strains under controlled greenhouse conditions and assessed their agro-morphological performance under field conditions representative of local agroecological conditions. Greenhouse pathogen assays demonstrated that GE’d IR64 and Ciherang-Sub1 lines were resistant to all tested local Xoo strains across three successive generations. We identified TalC as the primary disease-causing effector in the local Xoo populations. Irrigated field trials conducted over two seasons in the Kou Valley, Burkina Faso, revealed absence of agro-morphological penalties in GE’d lines compared to their parental wild-type lines. Observed trait variation was attributable to environmental fluctuations rather than genomic modifications. Collectively, our findings demonstrate that genome editing of the rice lines does not impose growth penalties, and support the suitability of GE’d IR64 and Ciherang-Sub1 for large-scale adoption in Burkina Faso, pending multi-location validation and introgression into locally adapted varieties.

## Introduction

Rice is a strategic crop in Burkina Faso, ranking first among imported cereals and fourth among cultivated cereals by area and production. In Burkina Faso, rice consumption is increasing at an annual rate of 7 %, driven by population growth, rapid urbanization, and evolving dietary habits. The current national demand is estimated at approximately 900,000 tons per year, whereas the average national production in 2023 was only 50,424 tons^1^. To bridge this gap, Burkina Faso relies heavily on rice imports, which cost over 40 billion CFA francs annually^2^. In response to this import dependency, the government of Burkina Faso has prioritized efforts towards enhancing local rice production. However, these efforts are hindered by persistently low yields, averaging at just 2.2 tons per hectare (https://www.fao.org/home/en). This low productivity is largely attributed to various abiotic and biotic constraints that limit the performance of cultivated rice varieties.

One of the most significant biotic constraints to rice production in Burkina Faso is bacterial leaf blight (BB), caused by *Xanthomonas oryzae* pv. *oryzae* (Xoo). BB can affect rice at any growth stage but is most prevalent at maximum tillering and maturity, where it induces lesions that can lead to wilting and, in severe cases, complete plant death^3^. BB was first reported in Burkina Faso in 1981, and major outbreaks were recorded in 1998 and 2004, particularly in the Bagré rice-growing plains, resulting in yield losses exceeding 50%^4^. More recently, incidence rates exceeding 80% have been reported in several rice production areas^5^. It is particularly concerning that the majority of irrigated and lowland rice varieties currently grown in Burkina Faso are highly susceptible to BB^6^. Since the irrigated lowland cultivation system contributes more than 90% of the national rice supply, there is an urgent need for effective strategies to enhance the resistance of cultivated rice varieties against BB to ensure sustainable rice production.

While a large number of BB resistance genes are known, individual R genes are often insufficient for effective host plant protection against BB and can be overcome by newly evolved or imported Xoo strains^7, 8^. Notably, two recent outbreaks of BB in East Africa, caused by strains introduced from Asia, are spreading rapidly^8^. Xoo is a xylem-borne pathogen that injects Transcription-Activator-like effector proteins (TALe) virulence protein into host cells. The repeat variables di-residue (RVD) of TALes target specific promoter sequences (effector binding elements, EBE) to directly activate SWEET sucrose uniporter gene in the xylem parenchyma^9, 10^. Activation of one of three SWEETs, i.e., *SWEET11a, SWEET13*, or *SWEET14* is sufficient to cause host susceptibility to Xoo. Modification of four EBEs in the promoters of *SWEET11a, SWEET13*, and *SWEET14* by genome editing in the *japonica* variety Kitaake led to resistance to ∼100 diverse strains collected in Asia and Africa^11^.

IR64 was a dominant rice variety in several Asian and African nations. Due to its popularity with farmers, IR64 has been widely used as a parent in rice breeding programs. One important breeding outcome from IR64 was Ciherang, which shares similar grain quality and genetics with IR64 but offers improved yield^12^. Using CRISPR-Cas9 and Cpf1, five EBEs were successfully edited in IR64 and Ciherang-Sub1 to confer broad-spectrum resistance to diverse *Xoo* strains^11, 13^. Multi-year, multi-condition experimental field plots with the genome-edited (GE’d) IR64 and Ciherang-Sub lines in two geographical locations (in the absence of disease pressure) indicated that the GE’d lines behaved similarly to their respective wild-type (WT) parents^14^. The potential of GE’d IR64 and Ciherang-Sub1 GE’d lines promises a solution to the emerging problems in Burkina Faso.

In this study, we tested the GE’d IR64 and Ciherang-Sub1 lines against a set of five Burkinabe Xoo strains across three generations, and evaluated the agro-morphological performance of the GE’d lines over two trial periods under agroecological conditions in Burkina Faso. We report resistance of GE’d lines from both rice cultivars to local Burkinabe strains. Using the GE’d lines, we establish that the five Xoo strains tested, whilst possessing TalF, rely on TalC to induce susceptibility in WT rice lines. We demonstrate that the GE’d lines do not impose an agro-morphological trade-off and that climatic fluctuations account for the variation in seasonal agro-morphological traits. This study laid the groundwork for the suitability of the GE’d IR64 and Ciherang-Sub1 lines for large-scale adoption in Burkina Faso and neighboring countries.

## Materials and Methods

### Plant materials

The study involved seven *Oryza sativa* subspecies *indica* genotypes, including five CRISPR-Cas9 GE’d lines and their respective WT parental varieties, IR64-WT and CS-WT. Four GE’d IR64 lines, and three GE’d Ciherang-Sub1 lines carry modifications in the EBE of PthXo1 (*SWEET11a*), PthXo2 (*SWEET13*), PthXo3, AvrXa7, and TalC (*SWEET14*, Table 1). Two local elite varieties, TS2 and FKR19, were included for comparison. The rice lines were evaluated over three generations under confined greenhouse conditions.

**Table 1:** WT and GE’d rice varieties used in this study.

| Genotype | Characteristic | Developed by |
| --- | --- | --- |
| **CS-WT** | Wild type (WT) | IRRI |
| **CS-1h** | Genome edited (GE'd) | 11, 13 |
| **CS-3b** | Genome edited (GE'd) | 11, 13 |
| **CS-6d** | Genome edited (GE'd) | 11, 13 |
| **IR64-WT** | Wild type (WT) | IRRI |
| **IR64-7a** | Genome edited (GE'd) | 11, 13 |
| **IR64-136a** | Genome edited (GE'd) | 11, 13 |
| **IR64-5d** | Genome edited (GE'd) | 11, 13 |
| **IR64-9a** | Genome edited (GE'd) | 11, 13 |
| **TS2** | Local susceptible variety (WT) | INERA |
| **FKR19** | Local resistant variety (WT) | INERA |

### *Xanthomonas oryzae* pv. *oryzae* strains used for pathogen test

Five *Xoo* strains, representing the genetic diversity of the strain^6, 15^, collected from various sites in Burkina Faso, were used to evaluate rice lines in Table 1. The characteristics of the strains are presented in Table 2. These strains are available in the collections at INERA and IRD Montpellier. They were collected from rice fields between 2003 and 2018, and isolated from leaf samples with BB symptoms. Xoo characterization was performed using Restriction Fragment Length Polymorphism (RFLP) markers and pathogenicity assays.

**Table 2:** List and features of Burkinabe strains used for virulence assays in this study.

| Strain | Race | Origin | Pathogen test |  |  |  |  | Major TALE | Reference |
| --- | --- | --- | --- | --- | --- | --- | --- | --- | --- |
|  |  |  | IR24 | IRBB3 | IRBB4 | IRBB5 | IRBB7 |  |  |
| BAI4 | A2 | Bagré | MS | R | R | R | R | TalF, TalC | <sup>15</sup> |
| BAI55 | A10 | Sourou | S | MS | MR | R | S | TalF, TalC | <sup>6</sup> |
| BAI111 | A3 | Bagré | R | R | R | R | R | TalC | <sup>6</sup> |
| BAI169 | A3 | Sourou | R | R | R | R | R | TalF, TalC | <sup>6</sup> |
| M3-4 | A3 | Mogtédo | R | R | R | R | R | TalF, TalC | <sup>6</sup> |

## Evaluation of GE’d lines towards Xoo in a confined greenhouse

### Experimental design

A simple non-randomized block consisting of 60 pots arranged in a tray (12 varieties × 5 strains) was employed. The seeds of each rice variety and line were disinfected and pre-germinated for 48-72 hours prior to transplanting into soil pots (5 L) with three seedlings per pot. Three replicates were prepared per rice line.

### Pathogen inoculation

3-week-old plants were infected with Xoo using the leaf-clipping method. Xoo grown on LPGA media were resuspended in sterile distilled water to a final OD600 of 0.2. The tips of the two youngest leaves of each plant were clipped with scissors dipped in the bacterial suspension. A total of 12 leaves were inoculated per strain and per variety/line. After inoculation, the plants were thoroughly watered for 48 hours to promote infection. Lesion length was measured 14-days post-infection.

### Evaluation of the agro-morphological performance of GE’d lines under confined field conditions Study site

This study was conducted in the irrigated rice-growing area of the Kou Valley, located in western Burkina Faso, approximately 30 km northwest of Bobo-Dioulasso, within the rural commune of Bama. The region has a South Sudanian climate, characterized by a distinct rainy season from mid-May to mid-October and a dry season from mid-October to mid-May. Annual rainfall is highly variable, averaging around 900 mm. The vegetation in the Kou Valley consists of shrub and tree savannah. The soils in the study area are hydromorphic with diverse textures.

### Field trial management

The soil in the experimental plot was tilled to a depth of approximately 25 cm using a power tiller. The experimental site was divided into plots prior to transplanting. Transplanting was carried out when the seedlings were 20 days old, with one seedling per hill, spaced at 20 cm × 20 cm both between rows and within rows. Water was supplied as needed. 200 kg/ha compound NPK fertilizer (14-23-14) was applied at transplanting. 150 kg/ha of urea (46% N) was applied in two split applications. 50 kg/ha was applied 15 days after transplanting, at the early tillering stage, while the remaining two-thirds were applied 55 days after transplanting, at the panicle initiation stage. Weed control was performed through manual weeding and harrowing as needed. No phytosanitary treatments were applied. At maturity, harvesting was done manually using sickles when two-thirds of the panicles had turned straw-colored. Each elementary plot was then threshed, winnowed, and dried.

### Agro-morphological traits evaluated

The agronomic traits were assessed according to the standard evaluation system scale of the International Rice Research Institute (IRRI)^16^. The following traits were measured:

1. Average number of tillers per hill at 60 days after transplanting (T60): Determined by manually counting the number of tillers on five randomly selected hills in each elementary plot.
2. Plant height at maturity (Height): Measured in centimeters (cm) at maturity on the tallest tiller using a graduated ruler from the soil surface to the tip of the panicle or the highest leaf. Measurements were taken on five randomly selected hills.
3. Number of panicles (Npan): Determined manually per hill before harvest.
4. Sowing-to-heading cycle (SHC): Defined as the number of days between sowing and 50% heading, recorded when 50% of the plants reached the heading stage.
5. Sowing-to-maturity cycle (SMC): Corresponds to the number of days between sowing and plant maturity when three-quarters of the panicles turned straw yellow.
6. Panicle weight (Wpan): Measured in grams (g) using a laboratory scale.
7. Panicle length (Lpan): Measured in cm at maturity after harvest using a graduated ruler.
8. Sterility rate (Sterility): Evaluated by counting the number of empty and filled grains on five panicles. The sterility rate was calculated using the formula: Sterility = (Number of empty grains/Total number of grains) x 100.
9. Thousand-grain weight (TGW): Weight in grams (g) of 1000 grains at a moisture content of 14 %.
10. Yield: Assessed in kg per elementary plot at a moisture content of 14% and extrapolated to a per-hectare basis.
11. Climatic data, in particular temperature, rainfall, and number of rainy days, were collected at the INERA research station in the Kou valley during the two years of the study (mid-May to mid-October 2023 and mid-October to mid-April 2024).

### Data processing and analysis

The field data were first recorded in a field notebook before being entered and processed using Excel spreadsheet software, version 2016. The XLSTAT software, version 2016, was used to perform normality tests. Agronomic trait data were analyzed using R. Statistical significance was assessed using Student’s *t*-tests and FDR. Linear mixed-effects models were fitted to account for experimental design and environmental variation, with genotype treated as a random effect and cultivar and editing status as fixed effects. Broad-sense heritability (H^2^) was estimated as the proportion of total phenotypic variance attributable to genetic variance. Statistical significance was determined at *P* < 0.05.

## Results

### GE’d lines are resistant to local Burkinabe Xoo strains under confined greenhouse conditions

Five representative local Burkinabe Xoo strains, i.e., BAI4, BAI55, BAI111, BAI169, and M3.4, were selected based on their geographic distribution and genetic diversity^6^. The virulence of the five strains was tested against five rice lines that carry individual R genes developed by IRRI (Table 2). While the control rice line IR24 was susceptible to BAI4 and BAI55, the lines carrying individual R genes IRBB3, IRBB4, IRBB5, and IRBB7 were resistant to all Xoo strains tested, except for BAI55, in which only IRBB5 was resistant (Table 2). To evaluate the virulence of the five Burkinabe strains (Table 2), leaves of three-week-old plants were clip-inoculated, and the disease symptoms, i.e., lesion length, were measured two weeks post-inoculation. Two local rice lines were included in the analysis, namely TS2 and FKR19. The local TS2 variety was highly susceptible, whereas FKR19 was resistant to all tested strains (Figure 1). The resistance of FKR19 to all tested strains is consistent with previous reports indicating stable resistance of FKR19 to local Xoo strains under greenhouse and field conditions^6, 17^.

**Figure 1:**
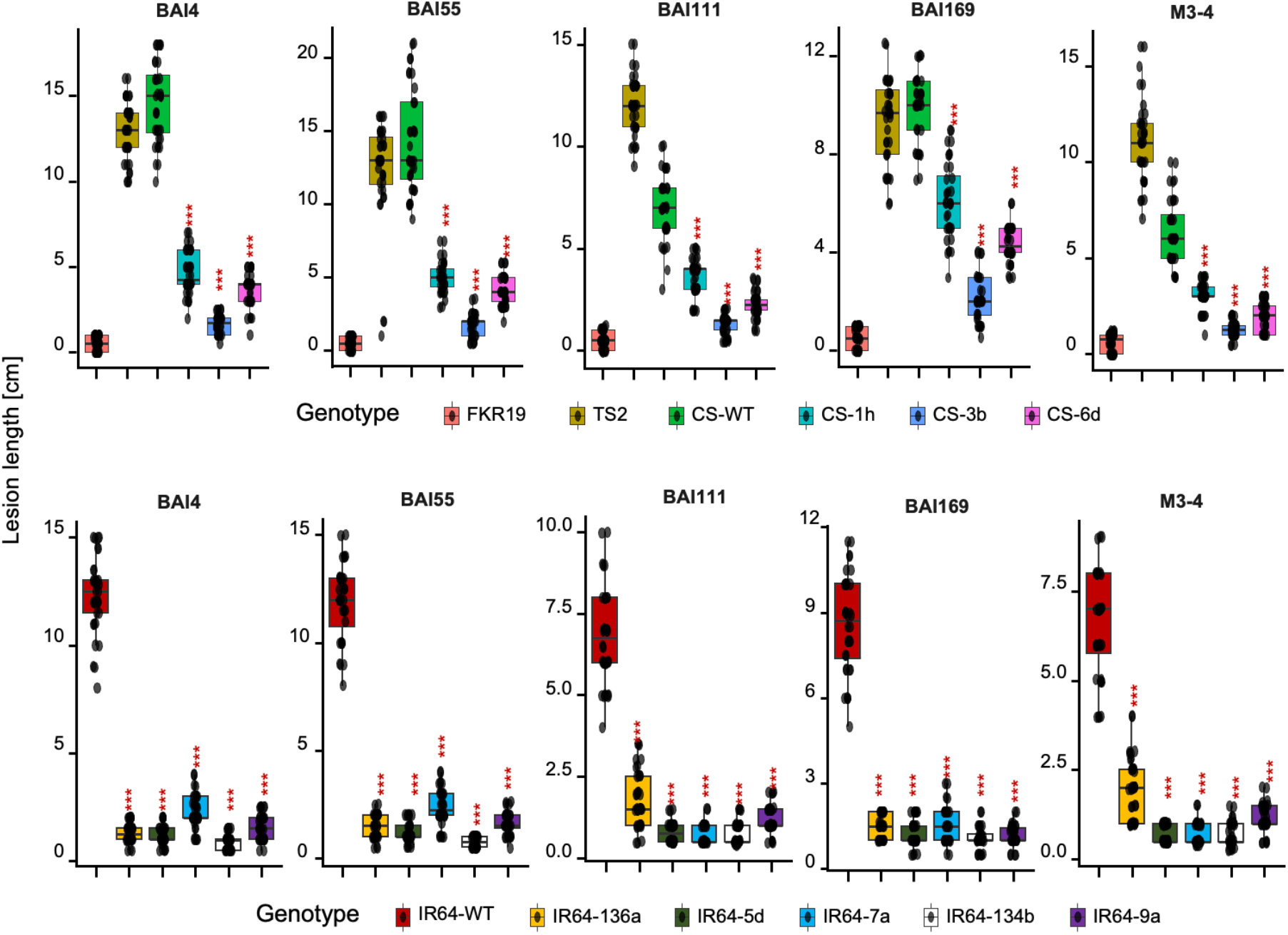
Lesion length quantification of WT and GE’d lines infected by local Burkinabe Xoo strains. week-old Ciherang-Sub1 (CS) or IR64 plants leaf clip-inoculated with five local Burkinabe Xoo strains. Lesion lengths were measured 2 weeks post-inoculation. Black dots indicate individual data points, the boxplot spans the interquartile range, horizontal line inside the box indicates the median, and the whiskers extend to the most extreme non-outlier values. Asterisks indicate statistical significance calculated using Student’s T-test relative to the respective WT. N > 30 from 3 independent repeats.

WT Ciherang-Sub1 and IR64 were susceptible to all five strains, as indicated by lesion development compared with the resistant cultivar FKR19 (Figure 1). Disease progression in WT Ciherang-Sub1 and IR64 plants could be attributed to the presence of the major TALes TalF and/or TalC in the tested Xoo strains^18^ (Table 2). Compared to the susceptible rice cultivar TS2, WT Ciherang-Sub1 and IR64 were less susceptible to BAI111, with a mean lesion length recorded for TS2 was 12.1 cm, whereas 7.1 cm and 6.8 cm for WT Ciherang-Sub1 and IR64, respectively; and Xoo strain M3-4 whereby the mean lesion length for TS2 was 11.1 cm, whereas 6.5 cm for both WT Ciherang-Sub1 and IR64 (Figure 1, Table S1). WT Ciherang-Sub1 and IR64 were equally susceptible as TS2 to the Xoo strains BAI4, BAI55, and BAI169, with a difference of less than 2 cm in mean lesion lengths.

Across three generations, all GE’d lines from both cultivars showed reduced susceptibility to the five Xoo strains tested, evident by significantly reduced lesion lengths compared to respective WT parental lines and susceptible cultivar TS2 (Figure 1, Table S1). Although the lesion lengths caused by BAI55 and BAI169 were more than 5 cm in the GE’d line CS-1h, the lesion lengths were 66% (BAI55) and 30% (BAI169) reduced in the CS-1h compared to WT Ciherang-Sub1. Intriguingly, although the GE’d lines retained WT EBE sequence for TalF binding (thus presenting susceptibility to BAI4, BAI55, BAI169, and M3-4), the GE’d lines were resistant to all the Xoo strains tested (Figures 1 and 2). The compatibility between the RVDs of two TalF variants (derived from BAI3 and MAI^19^) and the TalF EBE in IR64 and Ciherang-Sub1 was examined. Two variants of a TalF differ in their compatibility to bind to the EBE of *SWEET14*. For TalF derived from BAI3, the EBE contains two consecutive 2 bp mismatches followed by a 1-bp match, and an additional 4 bp mismatches, which likely reduces EBE binding affinity, thus preventing activation of *SWEET14* in IR64 and Ciherang-Sub1 (Figure 2). The resistant phenotype is consistent with the lack of predicted EBE in the upstream of *SWEET14* for TalF from BAI3^19^. In contrast, TalF from MAI1 contains only single-base mismatches interrupted by longer complementary regions (Figure 2). Further, TalF could undergo conformational changes, i.e., nucleotide looping out, to establish more efficient binding^20^ to the EBE upstream of *SWEET14*, thereby enabling disease development. Taken together, the data suggest that the Burkinabe strains carry TalF RVDs that resembles that of BAI3 rather than MAI1, therefore were unable to induce *SWEET14* for virulence. Importantly, the GE’d lines are not only resistant to Asian and East African strains^11, 13^, but also to the various Burkinabe strains.

**Figure 2:**
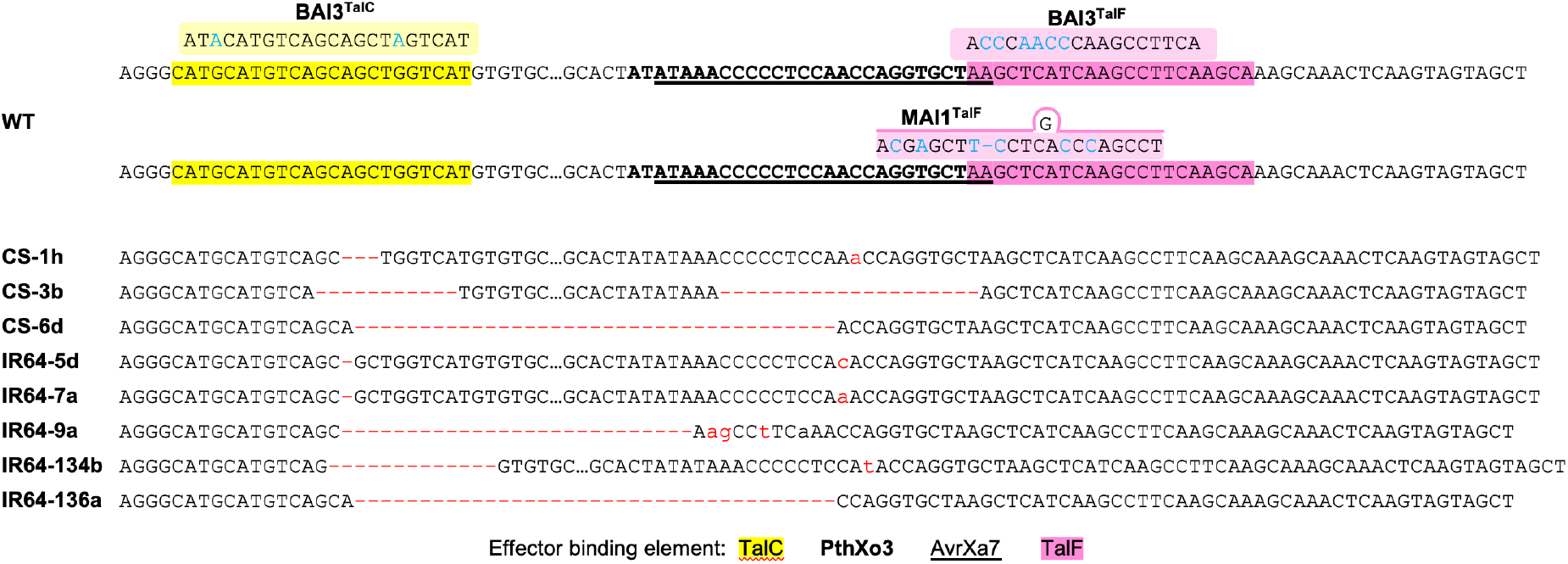
Effector binding elements (EBE) upstream of *SWEET14* in IR64 and Ciherang-Sub1. EBEs of WT (IR64 and Ciheran-Sub1) and respective GE’d IR64 and Ciherang-Sub1 lines, indicated with corresponding major TALe binding sites. Marked in red are edits introduced into the GE’d lines, in blue are mismatches on the TALe. Boxed in yellow and pink are TalC and TalF, respectively, from the indicated Xoo strains.

### Agro-morphological traits of GE’d lines are comparable to WT in field trials

To evaluate whether the GE’d lines could exhibit potential penalty in agronomic performance under field conditions in Burkina Faso, ten agronomic traits (Methods) were quantified from plants grown in the wet season of 2023 and the dry season of 2024 (Table S2). Principal component analysis (PCA) was performed to obtain an overview of the agro-morphological traits of GE’d and WT lines from both cultivars across the two trials. PCA resulted in four main clusters, each cluster consisting of WT and GE’d lines of each cultivar (Figure 3A). The clusters were separated by differences in cultivar (PC1-61.6% of the variation explained) and trial season (PC2-15.8% of the variation explained), indicating that potential differences in overall agronomic performance between WT and GE’d lines were not significant.

**Figure 3:**
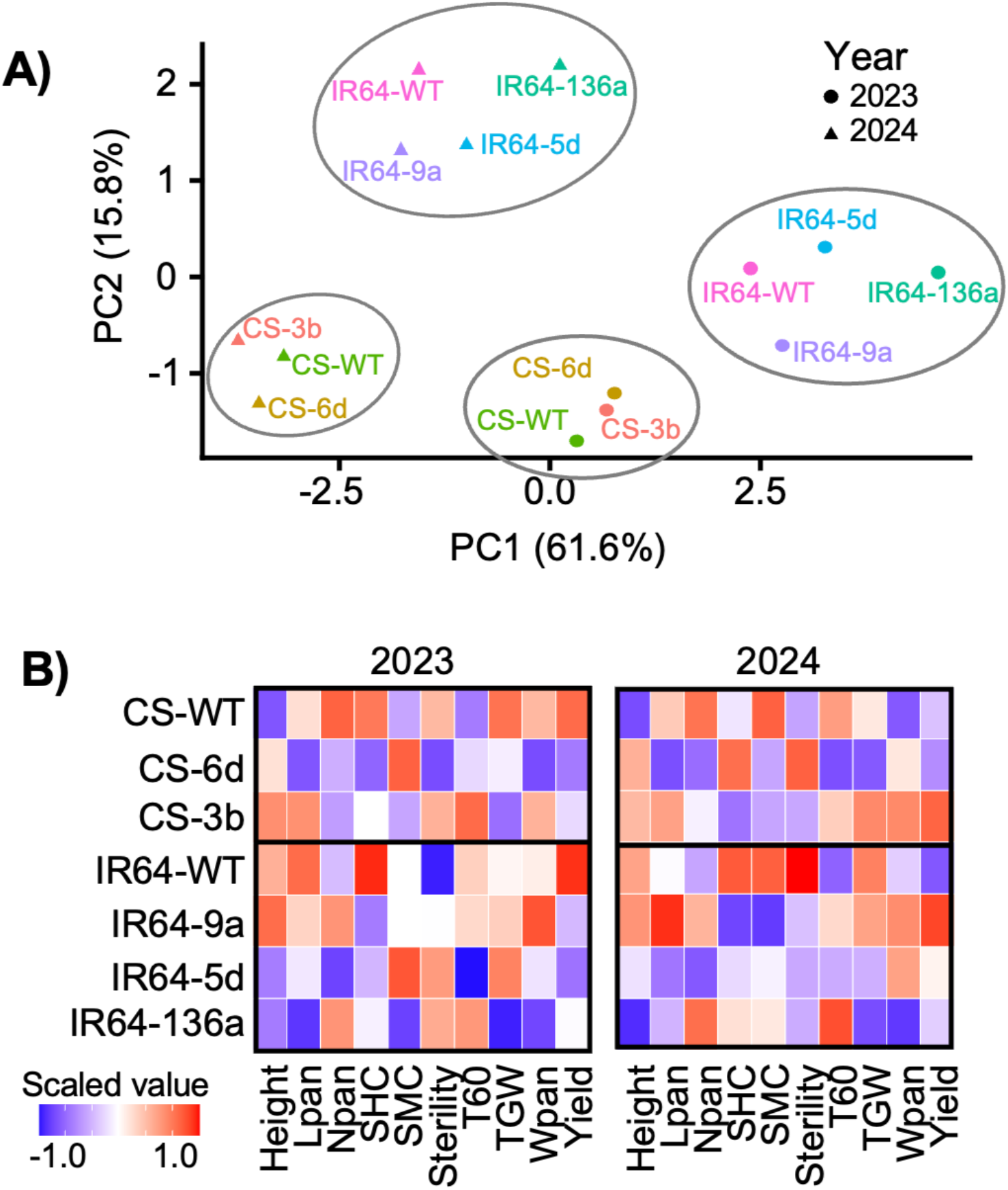
Agro-morphological traits of GE’d and WT lines across 2023 and 2024 experiment field trials. A) Principal component analysis on the 10 agro-morphological traits studied in field trials. Values in brackets of X- and Y-axes indicate variance explained for PC1 and PC2. Colors indicate genotypes, and shapes indicate trial season/year. B) Heatmap depicting quantitative agro-morphological traits standardized by Z-score normalization within each trait. Colour gradient indicate the scaled Z-scores.

To assess the potential differences in individual agronomic traits among the lines, Z-scores were calculated and normalized within each trait, cultivar, and year, for objective comparison of genotypes across the two trials (Figure 3B). Across both trials, the growth patterns for height, Lpan, Npan, and sterility were consistent between WT and GE’d Ciherang-Sub1 lines (Figure 3B). Similarly, patterns for height, Npan, SHC, sterility, and TGW were consistent between WT and GE’d IR64 lines across both trials. Trait variabilities were observed for SHC, SMC, T60, TGW, Wpan, and yield for Ciherang-Sub1, and in Lpan, SMC, T60, Wpan, and yield for IR64, with the coefficients of variation ranging between 0.6% and 22% (Table S2). Importantly, none of the traits exhibited by GE’d lines were significantly different compared to their respective parental WT lines. Collectively, PCA and individual trait inspections indicated the absence of agro-morphological differences attributable to genome editing.

### Variation in argo-morphological traits are attributed to environmental interaction

Three agro-morphological traits exhibited consistent patterns across all GE’d lines regardless of cultivar. The SHC and yield in 2023, and the SMC in 2024 of GE’d lines were lower compared to respective WT lines (Figure 3B). To examine whether the agro-morphological traits were genetically encoded or influenced by the trial season (wet season in 2024 vs. dry season in 2024), broad-sense heritability was computed. The board-sense heritability measures the ratio of phenotypic variance across all GE’d lines relative to environmental variations. All agro-morphological traits, including the observed consistent SHC, yield, and SMC patterns across GE’d lines, showed a heritability score of less than 0.3, indicating that environmental variation or experimental noise accounted for the trait variance (Figure 4A). Since the TGW has a heritability score of more than 0.3, suggesting potential genetically encoded heritability, the TGW for each GE’d line was compared to its respective WT. The TGW for each GE’d line, regardless of cultivar, was not significantly different from WT across both trial seasons (Figure 4B).

**Figure 4:**
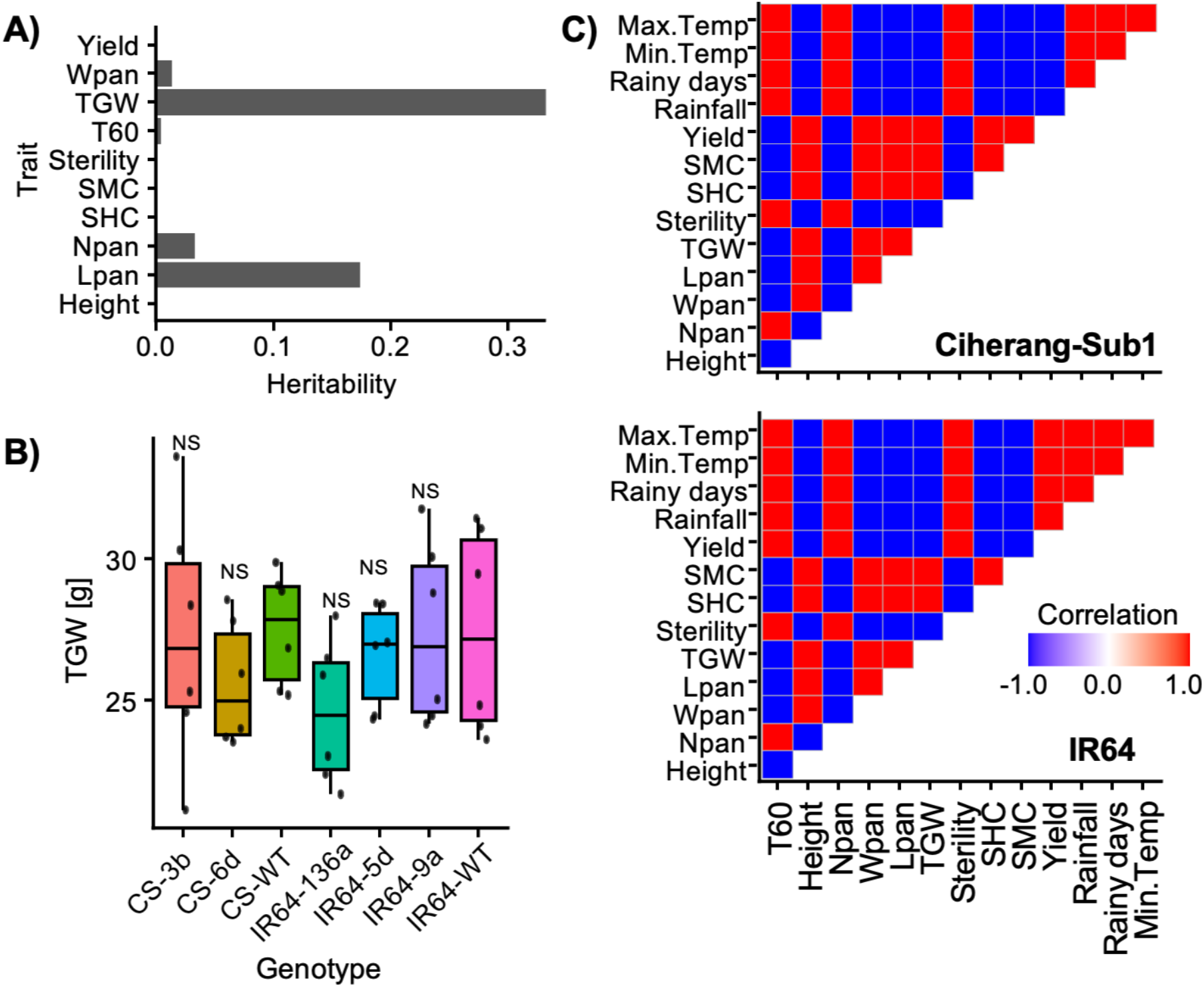
Heritability and correlation of agro-morphological traits and GE’d lines. A) Broad-sense heritability analysis for agro-morphological traits and environmental variables for all GE’d lines across 2023 - 2024. B) Thousand-grain weight (TGW) of each GE’d and WT line for Ciherang-Sub1 (CS) and IR64 lines. Statistical analysis performed with Student’s T-test for each GE’d line compared to their respective WT. NS: not significant C) Pearson correlation coefficients agro-morphological traits of GE’d lines and climate variables (excluding the genotypes CS-WT and IR64-WT). Data were aggregated by year and grouped according to cultivars CS (top) and IR64 (bottom). Color gradient indicates scaled Pearson correlation values.

To evaluate possible correlations among the agro-morphological traits and between traits and climate conditions, Pearson correlation coefficients were calculated. Positive correlations were observed between the T60 and Npan in the GE’d lines from both Ciherang-Sub1 and IR64 cultivars, suggesting that lines with more tillers tend to have higher panicle numbers, indicative of better vegetative growth and potential yield (Figure 4C). SHC and SMC were positively correlated with Wpan, suggesting that longer growth cycles may favor panicle development. However, negative correlations between T60 and both SHC and SMC suggest a trade-off between tillering and the duration of the growing cycle, potentially indicating an adaptive response to variable climatic conditions. Plant sterility showed positive correlations with rainfall and temperature, suggesting that excessive rainfall and elevated temperatures may negatively impact pollination, hence, grain formation. The correlations among agro-morphological traits and between climate conditions were consistent across GE’d lines of both cultivars, except for yield. In GE’d Ciherang-Sub1 lines, yield was positively correlated with plant height, Wpan, Lpan, TGW, SHC, SMC, but negatively correlated with T60, Npan, sterility, and all climate conditions tested. For GE’d IR64 lines, yield showed the inverse correlation observed for that of GE’d Ciherang-Sub1 lines (Figure 4C). Importantly however, the correlation analysis for GE’d lines was not different from their respective parental WT lines (Figure S1). In sum, trait x environment interaction studies indicated that variations in agro-morphological traits observed across two field trials were driven by environmental and climatic fluctuations rather than genome-editing.

## Discussion

### Effectiveness of modification of EBE for resistance towards local Burkinabe Xoo strains

We evaluated independent GE’d elite rice lines, i.e., editing the TALe binding sites (EBE) upstream of *SWEET11a, SWEET13*, and *SWEET14* (Figure 2) of IR64 and Ciherang-Sub1, against a panel of local strains from Burkina Faso. The results confirmed the efficacy of the strategy in conferring resistance to rice plants against five locally adapted Xoo strains (Figure 1). The resistant phenotype observed in Burkina Faso is consistent with previous reports^8, 11, 13^, in which the rice cultivars Kitaake, IR64, Ci-herang-Sub1, and Komboka, genome-edited using a similar strategy, exhibited broad-spectrum resistance against Asian and various African Xoo strains.

The time-resolved sampling and differences in TALome patterns support a model where Xoo populations in Burkina Faso are evolving under strong selection pressures. The IRBB lines were susceptible to older isolates, i.e., BAI4 and BAI55, but resistant to more recent isolates (i.e., BAI111, BAI169, M3-4), suggesting a shift in virulence strategies over time. Both host resistance deployment and environmental factors could contribute to the Xoo evolution, leading to the emergence and spread of new dominant races with altered virulence profiles. The dynamic evolution of Xoo highlights the need for more durable resistance strategies to anticipate Xoo adaptation and reduce the risk of future bacterial blight disease outbreaks.

Though BAI4, BAI55, BAI169, and M3-4 were reported to carry TalF^6^, the Restriction Fragment Length Polymorphism (RFLP) conducted was not able to precisely identify the TalF variants^19^. By comparing the EBE sequences upstream of *SWEET14*, this study concluded that the TalF in BAI4, BAI55, BAI169, and M3-4 could be a variant similar to that of BAI3, i.e., an inactive variant that cannot cause disease^19, 21^. Hence, the virulence of the Burkinabe strains tested in this study is highly likely attributed to TalC (Figure 2). In agreement, the loss of TalC EBE recognition in GE’d IR64 and Ci-herang-Sub1 lines resulted in resistance against the Xoo strains (Figures 1-2). Notably, caution is warranted, as the evolution of TalF through recombination and repeat reshuffling has been reported, which may confer an acquired ability to activate *SWEET14* and loss of host resistance^19^. Therefore, GE’d rice lines with disrupted TalF EBE^8^ may provide a more durable and future-proof resistance strategy against emerging Xoo.

All five Xoo strains tested carry TalB, previously shown to induce *TFX* and *ERF123* to trigger susceptibility in rice plants^19, 22^. IR64, Ciherang-Sub1, and Komboka carry *TFX* and *ERF123* with EBE sites complementary to TalB binding motif. However, editing the EBE of TalC was sufficient to confer resistance towards BAI3^8^, indicating that although TalB is present and can contribute to virulence, its effect targets other functions and is unable to fully compensate TalC induction of *SWEET14*. Hence, reinstating that TalC is the major functional driver of susceptibility in TalC-bearing Xoo.

### Agronomic performance analysis under local agroecological conditions

This study presents a significant contribution to the growing body of evidence supporting the agronomic equivalence of GE’d crops to their WT progenitors under real-world field conditions^14, 23, 24^. Specifically, we evaluated the performance of GE’d lines across multiple generations and growing seasons. Our results show the lack of agronomic trade-offs relative to the parental WT lines, indicating that genome editing did not disrupt the genetic or physiological networks underlying key agronomic traits (Figure 3). These findings are consistent with previous multi-location field trials conducted in Colombia and the Philippines, which assessed overlapping sets of GE’d IR64 and Ciherang-Sub1 lines and similarly reported no adverse effects on agronomic performance compared to WT parents^14^. Notably, two independent multi-location studies that spanned distinct geographic regions, seasons, agro-climatic conditions, and disease environments have converged on a consistent outcome, i.e., GE’d lines perform comparably to their WT parents. Collectively, our results substantially strengthen the evidence for agronomic equivalence in GE’d rice lines.

### Biosafety regulations and national dissemination of GE’d lines

To advance towards national deployment of the GE’d lines requires navigating regulatory and biosafety considerations. While regulations for the exemption of transgenic crops as conventional breeding material have not yet been established in Burkina Faso, a prerequisite for exemption in the growing number of countries with genome editing regulations is verification of the absence of foreign DNA. In severe cases, up to six Cas9 fragments were detected in GE’d rice line^14^, necessitating multiple generations of segregation and extensive molecular screening to identify transgene-free progeny. The import of GE’d IR64 and Ciherang-Sub1 lines without vector integration^14, 25^ into national breeding pipelines is pending the establishment of biosafety regulations for Burkina Faso.

## Conclusions

This study evaluated the resistance of GE’d IR64 and Ciherang-Sub1 rice lines and their parental varieties to the genetic diversity of Xoo strains from Burkina Faso under confined greenhouse conditions, and assessed agro-morphological performance in irrigated field trials under local climate. The GE’d lines, assessed over three generations, exhibited stable and enhanced resistance to a panel of five representative genetically diverse Burkinabe Xoo strains. Field evaluations conducted over two consecutive cropping seasons revealed a lack of agro-morphological penalties and that genome editing did not significantly alter agro-morphological traits compared with the parental varieties. Modifications of the EBEs upstream of *SWEET* susceptibility genes confer broad-spectrum resistance to Xoo derived from Asia, East Africa, and Burkina Faso. The GE’d lines are strong candidates for improving BB crisis in Burkina Faso without compromising agronomic performance.

## Author contributions

The conception of this study was led by WI. The methodology was developed by KS. Research activities were carried out by KS. Data analysis and interpretation were conducted by KS, WI, KKA, EL, WBF, BS, and BA. Resources were provided by EL, WBF, BS, WI, KKA, and BA. Data management was handled by KS, WI, KS, and EL. The manuscript was written by EL, WBF, WI, KKA, and BA. The overall supervision of the study was ensured by WI. All authors have read and approved the final version of the manuscript.

## Funding

This study was supported in part by the Bill & Melinda Gates Foundation to HHU, with a subcontract to the Institut de Recherche pour le Développement (IRD) and a second level subcontract to the Institut de l’Environnement et de Recherches Agricoles (INERA), by Deutsche Forschungsgemeinschaft (DFG, German Research Foundation) under Germany’s Excellence Strategy – EXC-2048/1 – project ID 390686111 (WF) and the Alexander von Humboldt Professorship to WBF.

## Data availability statement

Our FAIR data publication, prepared according to standard guidelines^26^ can be found under https://doi.org/10.60534/2w5tk-gea28

## Acknowledgments

The authors would like to express their gratitude the Healthy Crops Consortium, IRD, INERA and the École Doctorale Sciences et Technologies for their support in making this research possible. We thank DataPLANT and myFAIR, in particular Dr. Dominik Brilhaus, for data management and support.

## Conflicts of interest

The authors declare no conflicts of interest. The funders had no involvement in the design of the study, data collection, data analysis and interpretation, manuscript writing, or the decision to publish the results.

## Supplementary figures/tables

**Figure S1:**
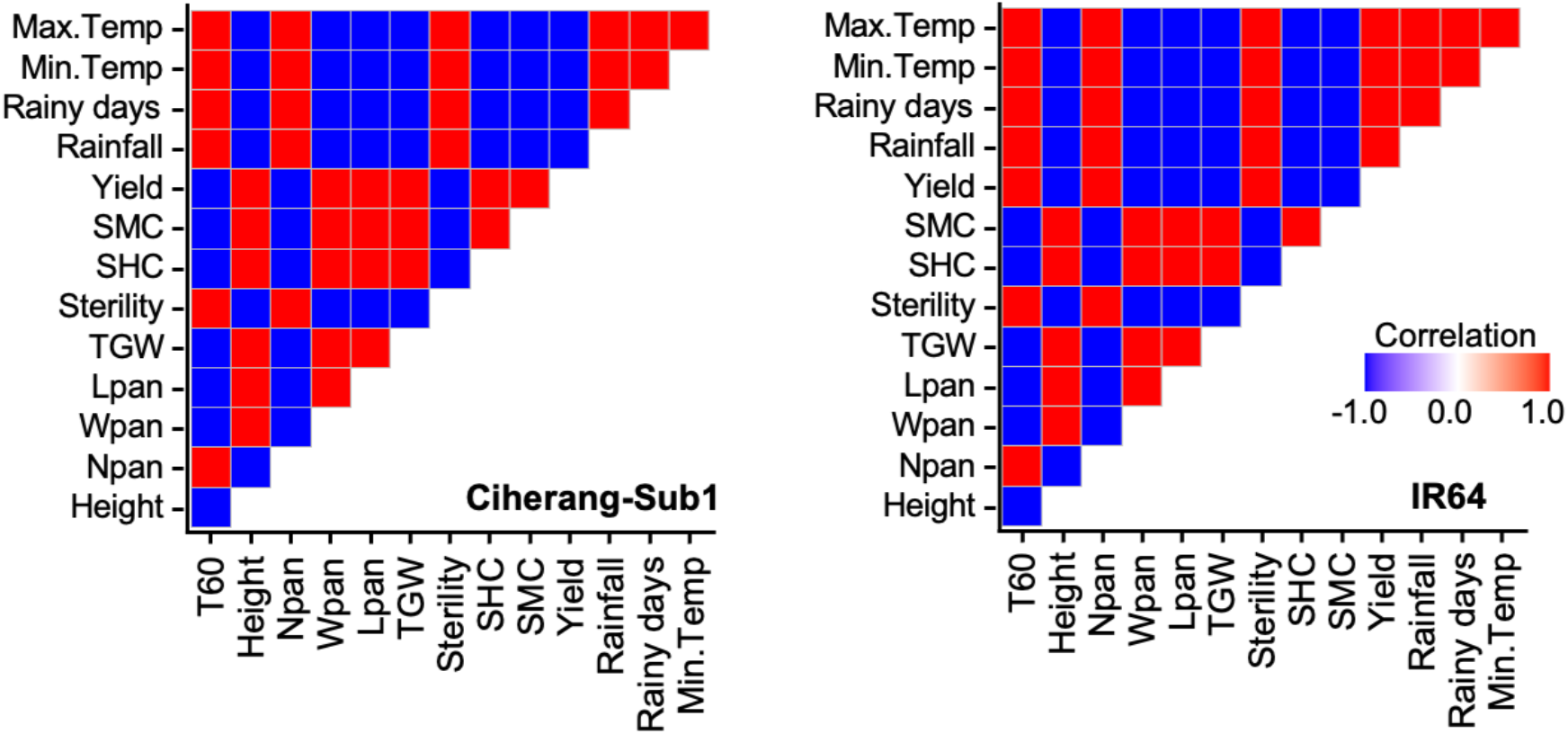
Correlation between agro-morphological traits of WT lines of each cultivar and climate across trial seasons. Pearson correlation coefficients agro-morphological traits of WT lines and climate variables. Data were aggregated by year and grouped according to cultivars CS (left) and IR64 (right). Color gradient indicates scaled Pearson correlation values.

**Table S1:** Lesion length quantification for WT and GE’d lines infected by local Xoo strains.

| Strain | Genotype | Mean [cm] | Replicate | Standard error | P-value |
| --- | --- | --- | --- | --- | --- |
| BAI111 | CS-WT | 7.06 | 36 | 0.27 |  |
| BAI111 | CS1h | 3.61 | 36 | 0.15 | 1.1 X 10<sup>-15</sup> |
| BAI111 | CS3b | 1.25 | 36 | 0.09 | 1.8 X 10<sup>-23</sup> |
| BAI111 | CS6d | 2.28 | 36 | 0.12 | 5.1 X 10<sup>-21</sup> |
| BAI169 | CS-WT | 9.89 | 36 | 0.23 |  |
| BAI169 | CS1h | 6.05 | 36 | 0.26 | 9.8 X 10<sup>-17</sup> |
| BAI169 | CS3b | 2.12 | 36 | 0.16 | 1.7 X 10<sup>-36</sup> |
| BAI169 | CS6d | 4.37 | 36 | 0.13 | 1.4 X 10<sup>-27</sup> |
| BAI4 | CS-WT | 14.59 | 36 | 0.39 |  |
| BAI4 | CS1h | 4.63 | 36 | 0.21 | 5.8 X 10<sup>-29</sup> |
| BAI4 | CS3b | 1.68 | 36 | 0.09 | 1.1 X 10<sup>-29</sup> |
| BAI4 | CS6d | 3.67 | 36 | 0.17 | 1.9 X 10<sup>-29</sup> |
| BAI55 | CS-WT | 14.22 | 36 | 0.58 |  |
| BAI55 | CS1h | 5.03 | 36 | 0.18 | 2.0 X 10<sup>-18</sup> |
| BAI55 | CS3b | 1.74 | 36 | 0.13 | 1.4 X 10<sup>-22</sup> |
| BAI55 | CS6d | 4.14 | 36 | 0.17 | 8.9 X 10<sup>-20</sup> |
| M3.4 | CS-WT | 6.47 | 36 | 0.28 |  |
| M3.4 | CS1h | 3.03 | 36 | 0.12 | 4.6 X 10<sup>-15</sup> |
| M3.4 | CS3b | 1.29 | 36 | 0.07 | 1.4 X 10<sup>-20</sup> |
| M3.4 | CS6d | 1.89 | 36 | 0.12 | 1.0 X 10<sup>-19</sup> |
| BAI111 | FKR19 | 0.51 | 36 | 0.07 | 5.4 X 10<sup>-26</sup> |
| BAI111 | IR64-WT | 6.81 | 36 | 0.25 |  |
| BAI111 | IR64134b | 0.76 | 36 | 0.06 | 8.5 X 10<sup>-25</sup> |
| BAI111 | IR64136a | 1.81 | 36 | 0.15 | 2.1 X 10<sup>-24</sup> |
| BAI111 | IR645d | 0.79 | 36 | 0.05 | 1.6 X 10<sup>-24</sup> |
| BAI111 | IR647a | 0.76 | 36 | 0.05 | 1.7 X 10<sup>-24</sup> |
| BAI111 | IR649a | 1.19 | 36 | 0.07 | 3.9 X 10<sup>-24</sup> |
| BAI111 | TS2 | 12.07 | 36 | 0.24 | 4.5 X 10<sup>-24</sup> |
| BAI169 | FKR19 | 0.54 | 36 | 0.07 | 2.3 X 10<sup>-27</sup> |
| BAI169 | IR64-WT | 8.55 | 36 | 0.28 |  |
| BAI169 | IR64134b | 1.07 | 36 | 0.06 | 7.6 X 10<sup>-26</sup> |
| BAI169 | IR64136a | 1.49 | 36 | 0.07 | 3.0 X 10<sup>-25</sup> |
| BAI169 | IR645d | 1.28 | 36 | 0.08 | 4.0 X 10<sup>-26</sup> |
| BAI169 | IR647a | 1.53 | 36 | 0.11 | 1.2 X 10<sup>-26</sup> |
| BAI169 | IR649a | 1.18 | 36 | 0.06 | 1.3 X 10<sup>-25</sup> |
| BAI169 | TS2 | 9.37 | 36 | 0.29 | 4.5 X 10<sup>-2</sup> |
| BAI4 | FKR19 | 0.48 | 36 | 0.07 | 3.1 X 10<sup>-33</sup> |
| BAI4 | IR64-WT | 12.32 | 36 | 0.29 |  |
| BAI4 | IR64134b | 0.84 | 36 | 0.06 | 1.7 X 10<sup>-32</sup> |
| BAI4 | IR64136a | 1.29 | 36 | 0.07 | 2.0 X 10<sup>-32</sup> |
| BAI4 | IR645d | 1.20 | 36 | 0.07 | 1.6 X 10<sup>-32</sup> |
| BAI4 | IR647a | 2.34 | 36 | 0.13 | 2.3 X 10<sup>-34</sup> |
| BAI4 | IR649a | 1.58 | 36 | 0.10 | 8.7 X 10<sup>-34</sup> |
| BAI4 | TS2 | 13.02 | 36 | 0.29 | 9.3 X 10<sup>-2</sup> |
| BAI55 | FKR19 | 0.47 | 36 | 0.07 | 2.5 X 10<sup>-32</sup> |
| BAI55 | IR64-WT | 11.92 | 36 | 0.29 |  |
| BAI55 | IR64134b | 0.75 | 36 | 0.04 | 6.4 X 10<sup>-31</sup> |
| BAI55 | IR64136a | 1.43 | 36 | 0.09 | 1.9 X 10<sup>-32</sup> |
| BAI55 | IR645d | 1.20 | 36 | 0.08 | 7.8 X 10<sup>-32</sup> |
| BAI55 | IR647a | 2.41 | 36 | 0.14 | 1.7 X 10<sup>-33</sup> |
| BAI55 | IR649a | 1.60 | 36 | 0.09 | 1.1 X 10<sup>-31</sup> |
| BAI55 | TS2 | 12.26 | 36 | 0.62 | 6.1 X 10<sup>-1</sup> |
| M3.4 | FKR19 | 0.61 | 36 | 0.08 | 1.9 X 10<sup>-25</sup> |
| M3.4 | IR64-WT | 6.56 | 36 | 0.25 |  |
| M3.4 | IR64134b | 0.72 | 36 | 0.06 | 3.1 X 10<sup>-24</sup> |
| M3.4 | IR64136a | 2.02 | 36 | 0.14 | 1.9 X 10<sup>-22</sup> |
| M3.4 | IR645d | 0.68 | 36 | 0.04 | 9.0 X 10<sup>-24</sup> |
| M3.4 | IR647a | 0.73 | 36 | 0.05 | 5.3 X 10<sup>-24</sup> |
| M3.4 | IR649a | 1.17 | 36 | 0.07 | 3.2 X 10<sup>-23</sup> |
| M3.4 | TS2 | 11.14 | 36 | 0.36 | 2.1 X 10<sup>-15</sup> |

**Table S2:**
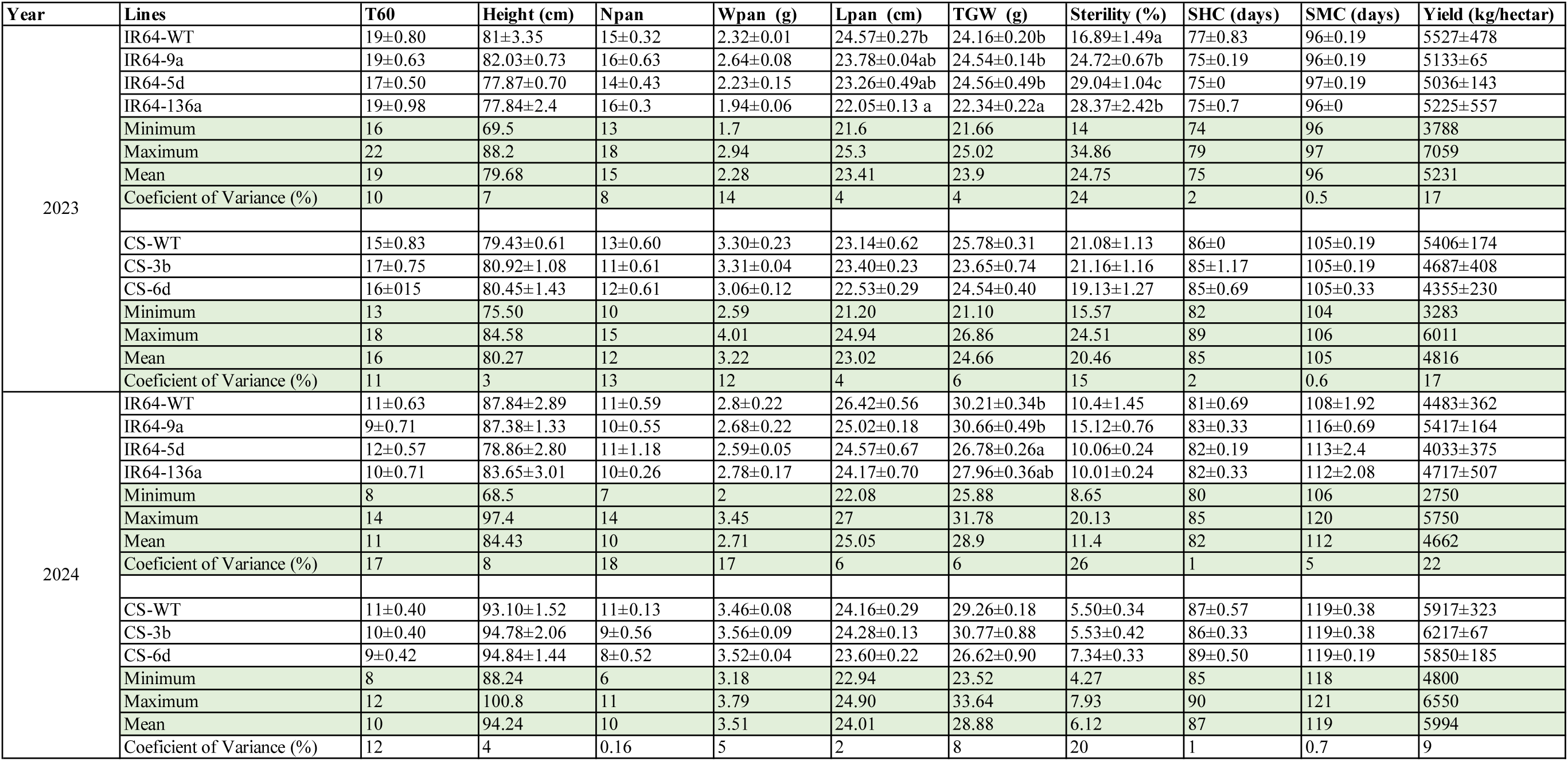
Agro-morphological traits of GE’d and WT lines in 2023 and 2024.

## Notes

### Competing Interest Statement

The authors have declared no competing interest.

